# Establishing thresholds for azole tolerance and persistence in *Aspergillus fumigatus t*o study their impact on voriconazole treatment *in vivo*

**DOI:** 10.64898/2025.12.26.693403

**Authors:** Rebeca Lobo-Vega, Diego Megías, Alba Río-Ropero, Álvaro Mato-López, Khalil Ashraph, Juliana Manosalva, Laura Alcázar Fuoli, Ana Alastruey-Izquierdo, Emilia Mellado, Jorge Amich

## Abstract

Antimicrobial tolerance and persistence are phenomena that enable pathogenic microbes to survive for extended periods in the presence of high concentrations of cidal drugs. Current evidence suggests that these phenomena may contribute to treatment failure and could even lead to the development of resistance. However, our understanding of antifungal tolerance and persistence in *Aspergillus fumigatus,* as well as their potential role in therapeutic failure, remains limited. In this study, we present an optimized, easy-to-perform method for detecting *A. fumigatus t*olerance and persistence to azole antifungals, based on a single colony-forming unit (CFU) measurement. Additionally, we developed a microscopic approach to investigate the dynamics of conidial killing in medium-throughput assays. Using these methods, we established epidemiological threshold values to classify strains as tolerant or persister and applied them to screen and categorize a collection of clinical isolates. Furthermore, we demonstrate that tolerance—but not persistence—negatively impacts the efficacy of voriconazole treatment in a *Galleria mellonella i*nfection model. Based on these findings, we propose that tolerance and persistence should be monitored in clinical isolates and potentially considered when determining therapeutic strategies.

## INTRODUCTION

*Aspergillus fumigatus i*s a saprophytic filamentous fungus that grows in soils feeding from organic decaying matter. During its natural life-cycle, this mold produces thousands of small spores or conidia that are distributed in the air to find new niches. Humans regularly inhale these conidia that, due to their small size, can reach deep in the respiratory tract [1]. This usually has no consequences for immunocompetent individuals, as conidia are very efficiently eliminated by the respiratory epithelium and the innate immune response [1]. Nevertheless, in immunocompromised patients and/or patients with underlying lung diseases, *A. fumigatus c*an cause lethal chronic and invasive infections [2]. Treatment of aspergillosis diseases is primarily based on the azole antifungals, with amphotericin-B and echinocandins as secondary or salvage lines [3–5]. Voriconazole (VCZ) and isavuconazole (IVZ) are generally employed as first line treatment for invasive aspergillosis [3–5]. For chronic aspergillosis, itraconazole or voriconazole are considered the best option for primary therapy [6]. Posaconazole is generally used as prophylaxis [7, 8]. Worryingly, in the last couple of decades azole resistance has spread globally [9], severely complicating the management of patients and increasing the mortality rates [10–12]. Besides the great relevance of azole resistance, it is important to remark that the mortality rates associated with invasive and chronic aspergilloses remain very high even in treated patients not infected with resistant isolates [13–19], demonstrating the importance to investigate other factors that may contribute to treatment failure.

Antimicrobial resistance is not the only mechanism by which pathogens can withstand the action of drugs. The phenomena of antimicrobial tolerance and persistence are gaining more attention in the last few years [20–23]. Both phenomena imply that pathogenic microbes can survive for extended periods in the presence of high concentrations (supra-MIC –minimum inhibitory concentration–) of cidal drugs. In tolerance all cells of an isogenic isolate can survive longer, whilst in persistence it is a sub-population cells that do so [20, 21]. Although still under scrutiny, the current evidence suggests that infection with tolerant or persister bacteria has a negative impact on the efficacy of antibiotic treatments [24–28].

In fungal pathogens most research has focused on the effects of fungistatic drugs, azoles for *Candida [*29–31] and echinocandins for *Aspergillus [*32, 33], but, little is known about tolerance and persistence to cidal drugs. We have previously shown that some *A. fumigatus i*solates can display persistence to voriconazole and isavuconazole [34], and suggested that azole persistence may contribute to treatment failure [35]. However, there is no data to delineate the thresholds that should be considered for tolerance or persistence detection, and therefore to evaluate the prevalence or the epidemiology of azole tolerance and persistence in *A. fumigatus s*trains. In this study we analyzed the prevalence of tolerance and persistence among all *A. fumigatus i*solates categorized as susceptible received in the Spanish Mycology Reference Laboratory in the years 2022 and 2023. To fulfil this task, we have optimized an amenable method to detect tolerance and persistence in the laboratory, and a microscopic pipeline to study the dynamics of killing in the presence of drug. Using these methods, we have established thresholds to determine persistence and tolerance in *A. fumigatus.* Moreover, we show that in the *Galleria mellonella m*odel of infection, tolerance, but not persistence, reduces the efficacy of a voriconazole treatment.

## RESULTS

### Azoles are fungicidal against Aspergillus fumigatus at high concentration

*A*ntimicrobial tolerance and persistence are defined as the capacity of some isolates to survive for extended periods in the presence of high (supra-MIC) concentrations of a cidal drug. Accordingly, in a recent perspective article we suggested that drug cidality should be confirmed before studying tolerance and persistence in filamentous fungi [36]. However, despite various publications showing high killing capacity, there is still reticence to classify azoles as cidal against *A. fumigatus,* because the stringent threshold of 99.9% killing in 24 hours generally employed for antibiotics [37] is not achieved. Nevertheless, as discussed in the perspective article, a less stringent threshold of ≥90% killing in 24 h (MDK90≤24h) has been proposed as a more appropriate criterion for filamentous fungi [36]. Therefore, to confirm the action of each of the three main families of antifungal drugs on *A. fumigatus,* we incubated 2.5×10^4^ conidia of various strains in the presence of high concentrations of voriconazole (VCZ, 8 µg/mL) amphotericin B (AMB, 16 µg/mL) or caspofungin (CSP, 8 µg/mL) and plated to count colony forming units (CFUs) at 24 and 48 hours. As expected from a true fungistatic drug, we could not assay CFUs for CSP as we observed substantial hyphal growth even at 24 h post-inoculation (not shown). Both AMB and VCZ killed more than 90% of the conidia of all tested strains in 24 h, therefore, achieving a MDK90<24h (Fig. 1A). At 48 h, AMB killed ≥99% of the spores (two-log decrease) of all strains (Fig. 1A), whereas VCZ killed >99% of the conidia of 3/5 strains, >98% of one isolate (PD9) and >95% of the remaining one (PD259) (Fig. 1). According to these results, we propose that both AMB and VCZ can be considered cidal against *A. fumigatus,* although some isolates appear to survive longer in the presence of VCZ and thus tentatively being tolerant or persister to this drug.

**Figure 1.**
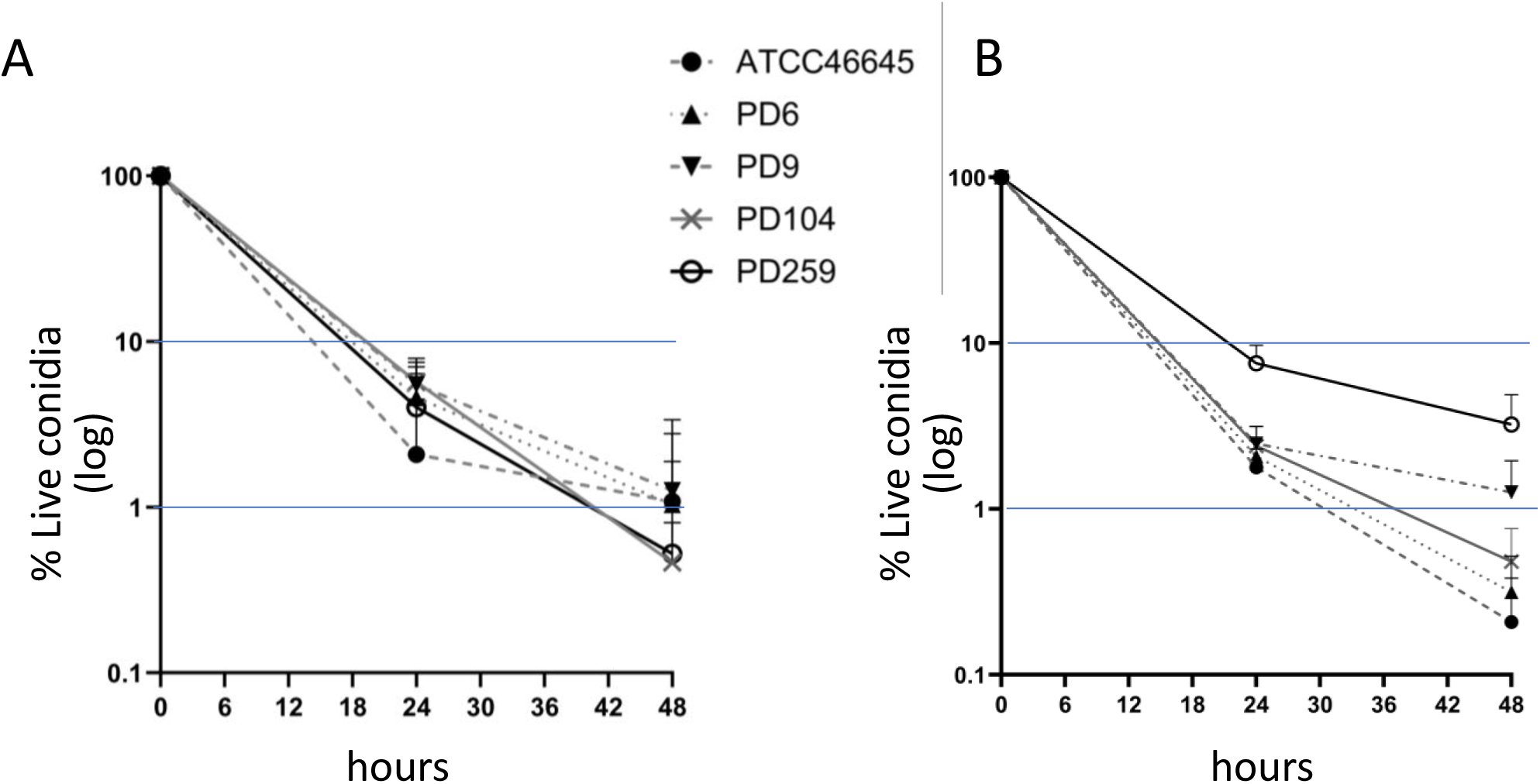
Amphotercin B and voriconazole are cidal at high concentration. Killing curves of five *A. fumigatus s*trains exposed to **A)** 16 µg/mL amphotericin B or **B)** 8 µg/mL voriconazole. Both drugs cause > 90% mortality to all strains in 24 hours. In other words, both drugs accomplished a MDK_90_<24 hours. Amphotericin B killing levels (**A**) were consistent for all strains, reaching MDK_99_ at around 48 hours. In contrast, voriconazole (**B**) displayed different levels of killing, with strains being highly killed (ATCC > 99.95%) and strains lowly killed (PD259 ∼95%) at 48 hours.

### Establishment of a single CFU-measurement method to detect azole tolerance/persistence

*S*ince tolerance and persistence are defined by the ability to survive for extended periods in the presence of supra-MIC concentrations of a cidal drug, the gold-standard method for their detection involves time-kill curve assays under high drug concentrations. [36]. These assays identify strains that exhibit prolonged survival compared to susceptible isolates. For instance, as can be observed in Fig. 1 the isolate PD259 is killed at a slower rate than the control strain ATCC46645, suggesting that it may be a persister or tolerant strain. However, since time-kill assays are labor-intensive, time-consuming, and prone to variability and technical errors when working with filamentous fungi, we consider them unsuitable for routine use in clinical laboratories. To address this limitation, we aimed to develop an alternative method to assess tolerance and persistence using a single CFU measurement at the endpoint of incubation. This approach was adapted from the EUCAST protocol for antifungal susceptibility testing. We inoculated an exact number of conidia (2.5×10^4^) in 96-well plates containing 8 µg/mL of the correspondent azole. After incubation the plate was centrifuged, the supernatant media (containing drug) discarded, the wells resuspended in 200 µL of NaCl-Tween20, thoroughly mixed and scraped, and the full content was plated onto rich potato dextrose agar (PDA) media plates. Preliminary experiments revealed that at 48 hours at 35°C there was high variance in the number of CFUs, which we believe may be due to the asynchronous germination of *A. fumigatus c*onidia. Therefore, we decided to extend the incubation to 72 hours, to ensure very high levels of conidial killing. With this longer incubation time we indeed observed very low numbers of surviving spores, resulting in a very narrow range of CFUs to detect different mortality rates. For instance, 99.9% killing would be detected with 25 CFUs and 99.95% with 12.5 CFUs. Consequently, we decided to assay a 10-fold higher inoculum (2.5×10^5^ conidia per well) to increase the range of countable CFUs. Using this optimized method (Fig. 2A), we could detect different percentages of survival after incubation with voriconazole (Fig. 2B) and isavuconazole (Fig. 2C). Therefore, this method can detect strains that survive longer in the presence of a high concentration of azoles with a single end-point measurement.

**Figure 2.**
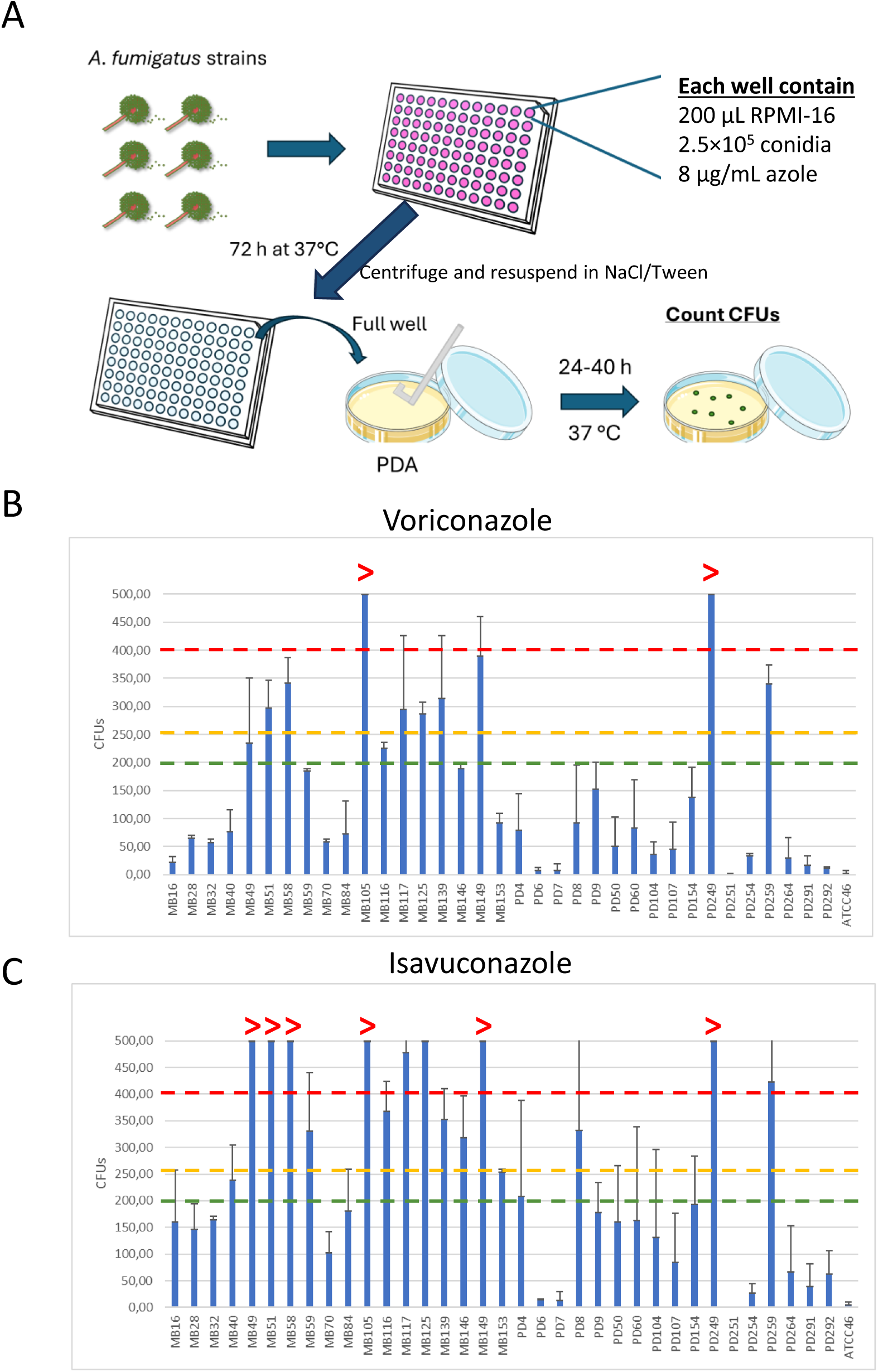
A single CFU-measurement method can detect different levels of survival upon exposure to high azole concentrations. A) Depiction of the method developed to detect different levels of survival in the presence of 8 µg/mL azoles with only one single-CFU measurement at final incubation time. The method was employed with a collection of 36 isolates, reflecting different levels of survival to **B)** voriconazole and **C)** isavuconazole among strains. The dashed lines mark the thresholds established later in study: susceptible below green line, undetermined between green and yellow lines, non-wild-type above yellow line (persister between yellow and red lines and tolerant above red line).

During the optimization of this method, we assayed four azoles voriconazole (VCZ), isavuconazole (IVZ), itraconazole (ITZ) and posaconazole (PSZ). We corroborated that this method was effective to detect different levels of survival for VCZ, IVZ and PSZ. However, it did not work for ITZ, where we always observed very high levels of survival after incubation with the drug. A close inspection revealed that at high concentrations, ITZ formed precipitation crystals. This observation suggests a local depletion of the drug next to the crystals, which allowed conidia in close proximity to these crystals to germinate and start growing (Fig. S1).

### Determination of the epidemiological thresholds for voriconazole and isavuconazole tolerance and persistence in A. fumigatus

*T*o gather epidemiological information of tolerance and persistence, we analyzed all the not resistant *A. fumigatus i*solates received in the Spanish Mycology Reference Laboratory in 2022 (=92, Table S1), 2023 (=55, Table S1), plus a collection of isolates previously used in the laboratory (=36, [34]) and using the laboratory strains ATCC44645 and Af293 as controls of high (>99.99%) and low (<99.8%) killing, respectively (as determined in preliminary experiments). We assayed the described method with 8 µg/mL of VCZ or IVZ, and we calculated the percentage of killing for each strain (counted CFUs / total inoculum). The methods to establish epidemiological cutoff values (ECOFF) for antifungal susceptibility are well defined [38]. However, it is important to note that these methods require the standard two-fold dilution series used in broth dilution assays to determine the MIC. This is because the ECOFF is calculated as the upper-end of the log-normal distribution derived from the MICs of a number of strains along the power of 2 dilutions. Although the type of data for survival is different, we decided to use ECOFFinder [39] to establish objective thresholds for non-wild-type behavior. We classified the strains in groups of discrete increasing numbers of counted CFUs, to mimic to some extent the classification of strains based on the MIC. The ECOFFinder results for voriconazole (Fig. 3A) and isavuconazole (Fig. 3B) suggested for both drugs a cut-off value of 200 CFUs (=99.92% killing) for non-wild-type behavior. Given the potential variability in CFU counts and the valley observed for 250 colonies (=99.90% killing) we considered prudent to denote that region as “undetermined” (Fig. 3A & 3B). More than 250 CFUs (<99.90% killing) would be established as non-wild-type. Interestingly, for both drugs two independent peaks can be observed in the non-wild-type population. We reasoned that they might be reflecting two different behaviors, thus tentatively classified them as persistence (from 250 to 400 CFUs, 99.90 to 99.84% killing) and tolerance (>400 CFUs, <99.84% killing) (Fig. 3A & 3B).

**Figure 3.**
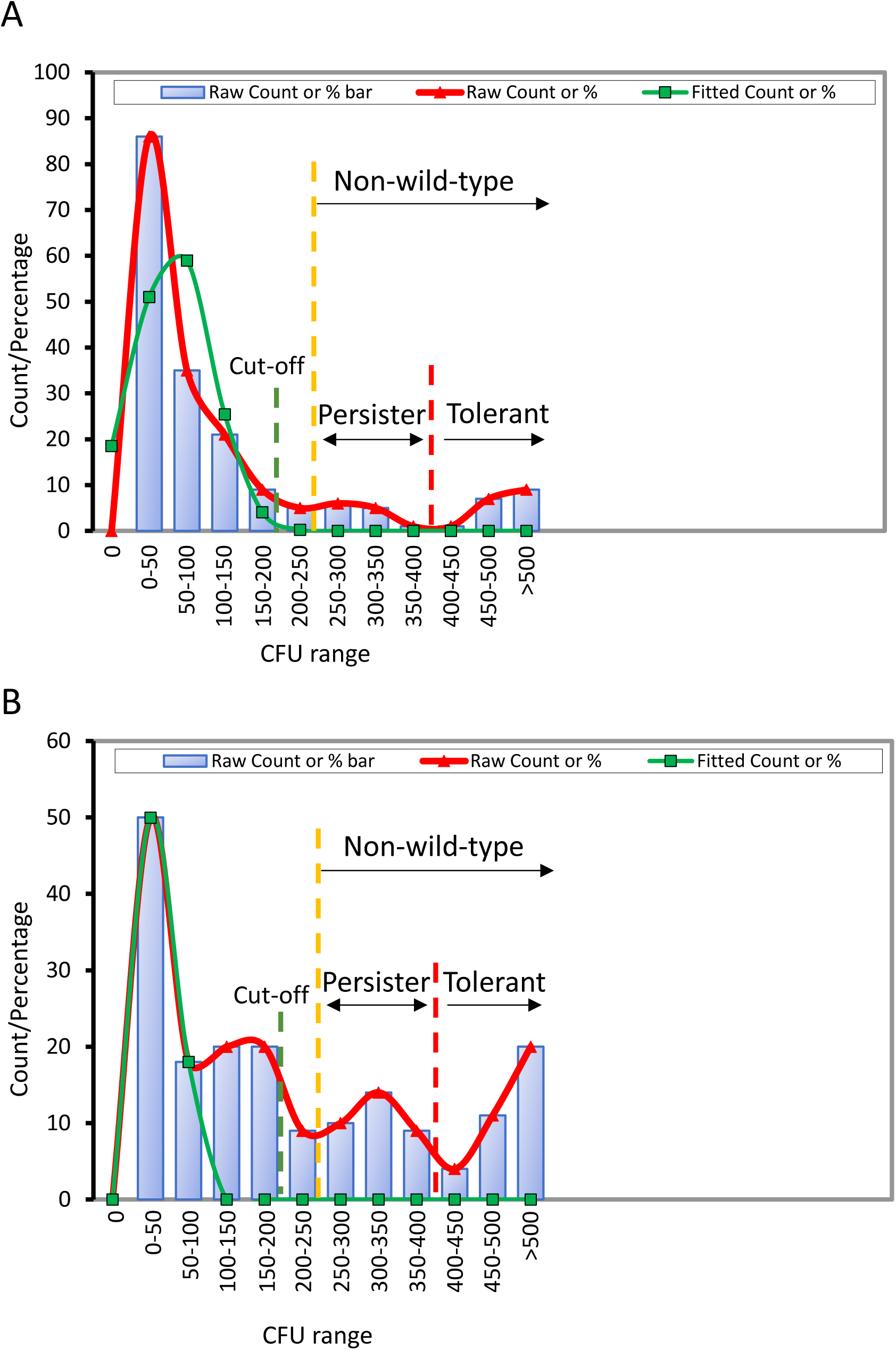
Determination of threshold values for azole tolerance and persistence in Aspergillus fumigatus using ECOFFinder. ECOFFinder provided a cut-off value of 200 CFUs (=99.92% killing) for both **A**) VCZ and **B**) IVZ (green dashed line). From 200 to 250 CFUs (99.2 to 99.90% killing) was categorized as “undetermined” (from green to yellow dashed lines). More than 250 CFUs (<99.90% killing) was categorized as non-wild-type behavior. Tentatively, from 250 to 400 CFUs (99.90 to 99.84% killing) was categorized as persister and more than 400 CFUs (<99.84% killing) as tolerant.

Using these thresholds, we could classify our collection of 183 *A. fumigatus s*trains into the defined groups (Fig. 4). We identified that 27/183 (14.8%) strains showed non-wild-type killing to VCZ and 68/183 (37.2%) to IVZ (Fig. 4A). Considering the tentative distinction between tolerance and persistence, we detected 11/183 (6.0%) strains displaying persistence and 16/183 (8.7%) tolerance to VCZ, and 35/183 (19.1%) of strains displaying persistence and 33/183 (18.0%) tolerance to IVZ (Fig. 4B). Therefore, it seems that persistence and tolerance are more common to IVZ than to VCZ. Indeed, whereas no strains were found to be tolerant or persistent solely to voriconazole, 41 strains showed non-wild-type behavior (persistence or tolerance) exclusively to isavuconazole.

**Figure 4.**
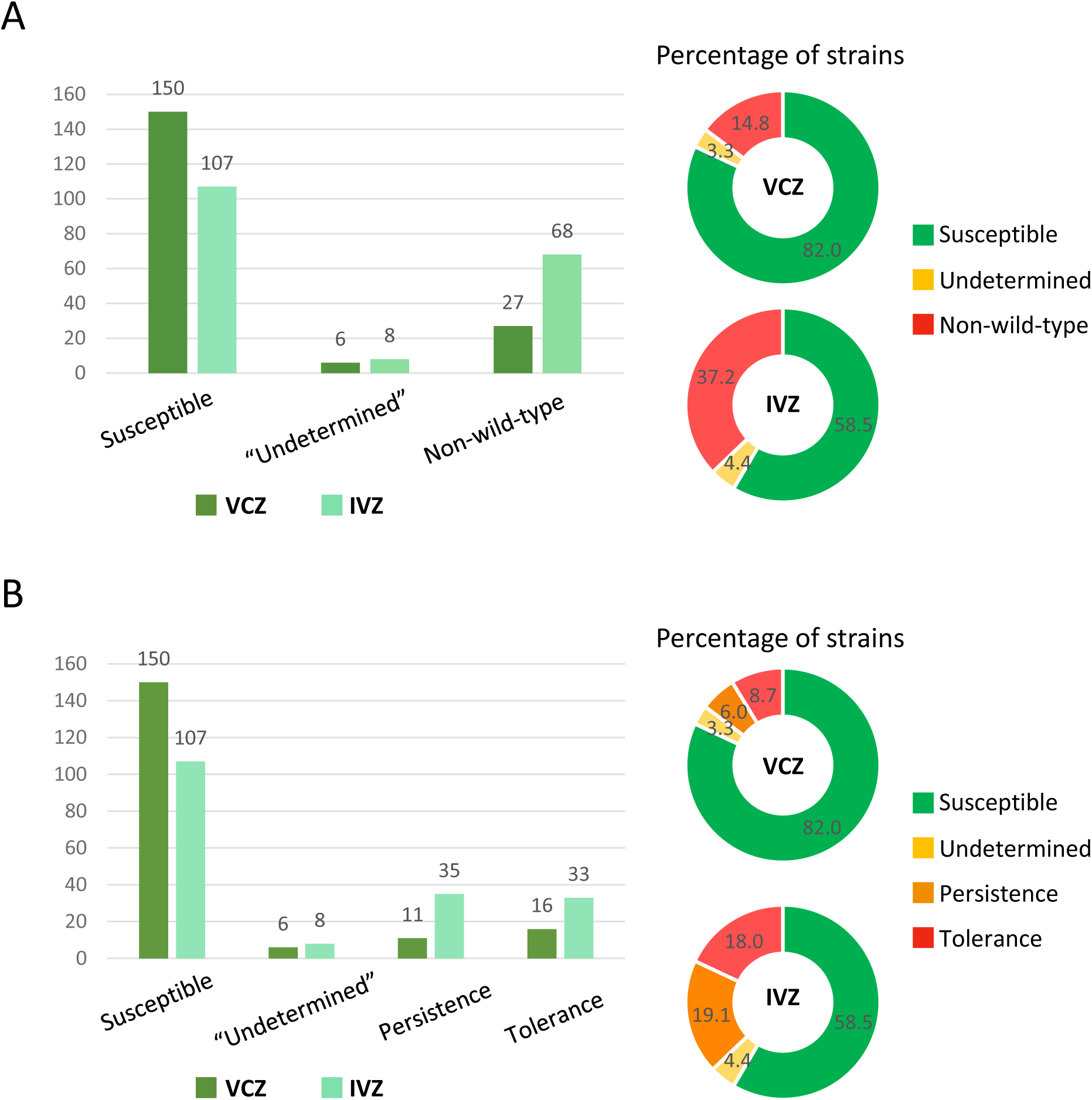
Classification of Aspergillus fumigatus strains using the single CFU measurement method. **A**) Distribution and percentage of survival behavior to voriconazole (VCZ) and isavuconazole (IVZ) in our collection of strains according to the thresholds established with ECOFFinder. **B)** Distribution and percentage of persistence and tolerance to voriconazole (VCZ) and isavuconazole (IVZ) in our collection of strains according to our tentative threshold classification.

### Establishment of a microscopy assay to investigate the dynamics of killing

To determine whether the survival levels detected using a single CFU measurement after incubation correlated with killing dynamics and given the challenges of performing CFU-based killing curves with filamentous fungi, we developed a microscopy-based method to monitor conidial death in real time. Additionally, we optimized an analytical pipeline to automatically quantify killing dynamics (see *Materials and Methods)*. Briefly, single conidia are followed during a period of 48 hours, and conidial death is defined as the time of Sytox™ signal activation in drug-exposed conidia.

Using this approach, we analyzed the killing dynamics of fourteen selected isolates (four susceptible, five persister, and five tolerant according to the previous classification) under 8 µg/mL of VOR or IVZ. The strains exhibited distinct killing patterns, clustering in agreement with their prior classification (Fig. 5A & B). Visually it can be observed that conidia of the susceptible strains initiated death earlier than other groups. The persister isolates began dying slightly later than the susceptible strains, with several reaching intermediate killing levels. Tolerant strains showed the longest delay before death onset and most exhibited lower killing levels. To assess the statistical significance of these observations, we quantified survival rate as the inverse of the initial killing slope (1/slope calculated over the first 10 hours of incubation). In this metric, higher values correspond to slower killing kinetics and therefore greater survival. In the presence of voriconazole, the survival rates of susceptible isolates ranged from 1.219 to 2.083, whereas those of persister isolates ranged from 2.945 to 5.249, and tolerant isolates ranged from 9.915 to 50.17 (Table S2). These differences were statistically significant (Fig. 5C). Similar trends were observed with isavuconazole, with survival rates differing significantly across groups (Table S2, Fig. 5D).

**Figure 5.**
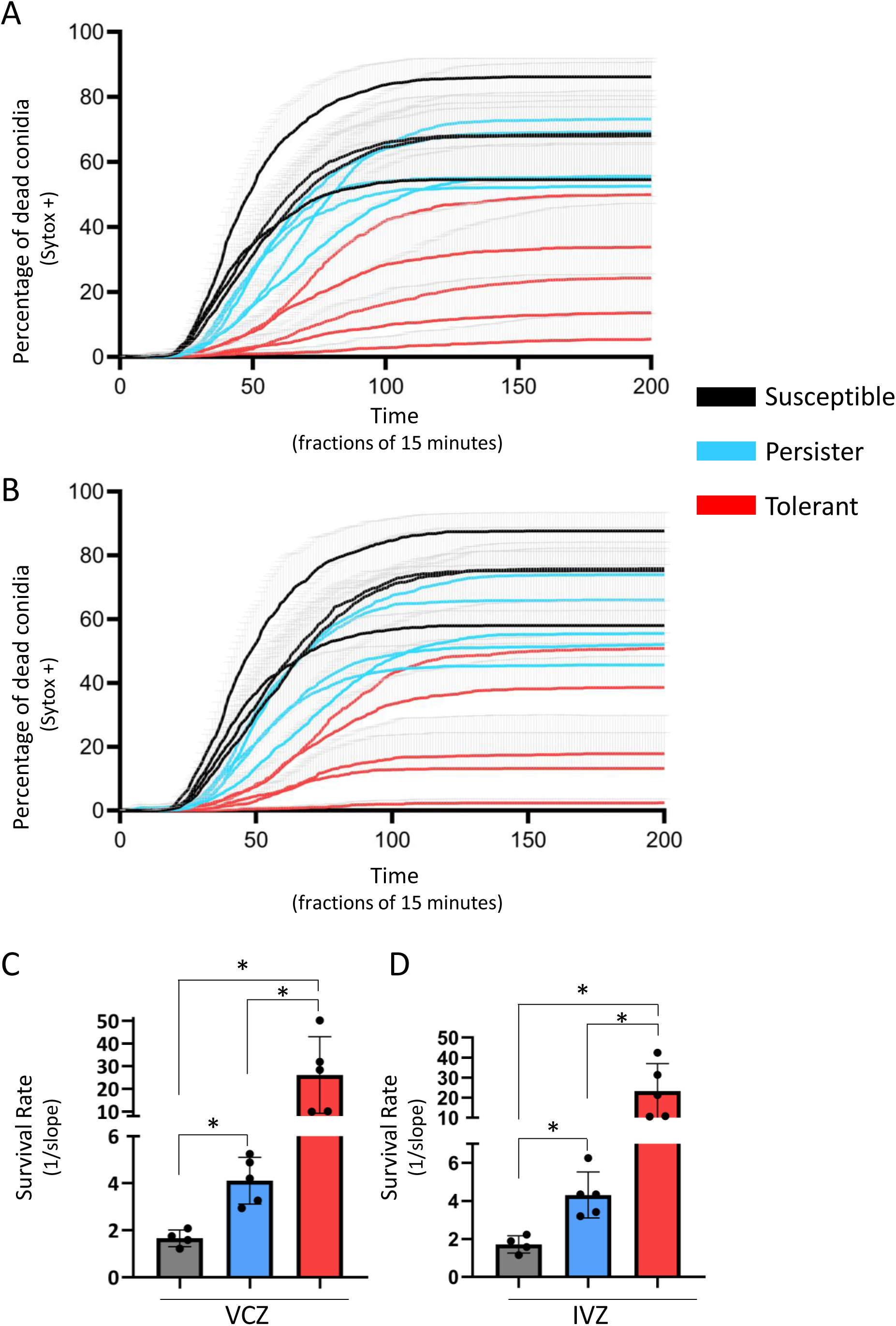
Killing curves of strains using the microscopy method display different dynamics of killing, which align with the results obtained with the single-CFU measurement. The killing dynamics of fourteen isolates, previously classified as susceptible (black: ATCC46645, CM-10565, CM-10773 and CM-1113), persister (blue: CM-10597, CM-10779, CM-10881, CM-10929 and CM-11429) or tolerant (red: Af293, CM-10571, CM-10677, CM-11185 and CM-11189) using the single-CFU measurement, were analyzed with the microscopy method upon exposure to **A)** voriconazole or **B)** isavuconazole. The susceptible isolates started dying first, followed by the persister isolates, and then the tolerant strains. The experiment was independently repeated four times. The inverse of the slope (1/slope) was calculated as a metric of survival for each strain, and the values compared among the three groups for **C)** voriconazole and **D)** isavuconazole. For both drugs, all groups showed statistically significant differences in survival rates. A Brown-Forsythe with Dunnett’s multiple comparisons was employed, *= *p-*value < 0.05,

Therefore, the killing dynamics captured by this methodology differentiated three different behaviors that corresponded to the previous classification, validating the reliability of the single-CFU measurement approach for detecting azole tolerance and persistence in *A. fumigatus*.

### Tolerance, but not persistence, decreases the efficacy of azole treatment in a model of infection

We previously showed that the fungal burden in *Galleria mellonella d*ecreased noticeably less after a voriconazole treatment in some of the larvae that had been infected with a persister isolate [34]. To further evaluate if azole tolerance or persistence may have an impact on the efficacy of treatment, and to test if the defined thresholds may serve to predict such negative effect, here we decided to investigate the efficacy of a voriconazole treatment against susceptible, persister or tolerant isolates in *G. mellonella*.

We infected larvae with 10^6^ conidia of susceptible (ATCC46645 and CM-10565) persister (CM-10597, CM-10767, CM-10779 and CM-10881) or tolerant (Af293, CM-10571, CM-10677, CM-11189 and CM-10909) strains, treated them with one injection of 4 µg/larvae of VCZ (equivalent to a therapeutic dose of 10 mg/kg/day [40]) at the time of infection, and monitored mortality for 7 days. The VCZ treatment had a significant beneficial effect on survival to larvae infected with all the susceptible (Fig. 6A) and persister (Fig. 6B) isolates. In contrast, the VCZ treatment did not provide a significant improvement on survival to larvae infected with 4 out of the 5 tested tolerant isolates (Fig. 6C).

**Figure 6.**
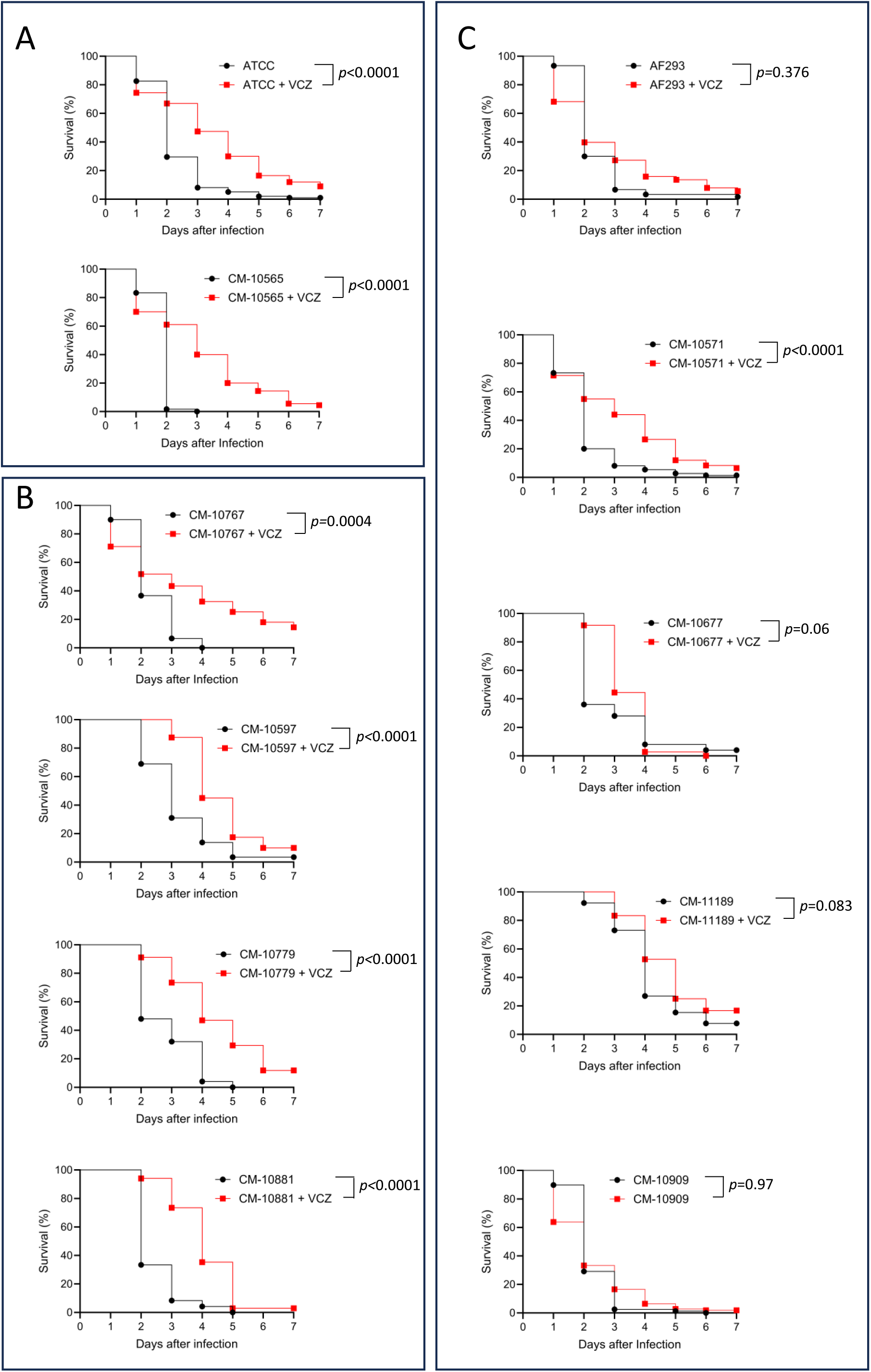
Infection with tolerant isolates cause a reduction in the efficacy of a voriconazole treatment. *Galleria mellonella l*arvae (groups of 20) were infected with 10^6^ conidia of susceptible (**A**), persister (**B**) or tolerant (**C**) isolates. The mortality caused by all susceptible and persister isolates was significantly reduced with a voriconazole treatment (4 µg/larva). In contrast, the voriconazole treatment did not provide a significant improvement in survival to larvae infected with four out of five tolerant isolates (Af293, 129, 137 and 354). The experiment was independently repeated two to four times.

Therefore, most of the tolerant strains did not respond as efficiently to a VCZ dose as susceptible isolates in this infection model, suggesting that tolerance may negatively impact the efficacy of treatment.

## DISCUSSION

The phenomena of tolerance and persistence are gaining relevance in fungal pathogens; however, the prevalence of these phenomena, as well as their potential relevance in treatment failure, are mostly unknown in *Aspergillus fumigatus*.

We recently published a perspective article where we proposed definitions and recommendations for investigating tolerance and persistence in filamentous fungi [36]. We advised that before investigating tolerance and persistence responses, the cidal capacity of the drug should be confirmed. To address this matter in our settings, we have conducted killing curve assays using the three main types of antifungal drugs. We have found that both amphotericin B and voriconazole kill >90% of spores within 24 h (MDK_90_ < 24 h), the threshold that we recommend to consider for drug cidality [36]. Therefore, in agreement with investigations showing cidal capacity of azoles against *A. fumigatus (*[41, 42], we suggest that azoles can be considered cidal against *A. fumigatus u*nder conditions of high drug exposure.

Interestingly, we observed some variability in the voriconazole killing dynamics for the different strains, whereas the killing achieved by amphotericin B was consistently very high. We speculate that this may reflect lower levels or even lack of tolerance/persistence to AMB in *A. fumigatus,* a hypothesis that deserves further investigations.

The gold-standard method to detect antifungal tolerance and persistence is the killing curve assay [36]. Nevertheless, this procedure is highly labor-intensive and error-prone, particularly when working with filamentous fungi. We consider it impractical to implement time-kill curve assays in clinical laboratories. Therefore, we have developed a simplified method for detecting extended survival that requires only a single CFU measurement at the endpoint of incubation. Using this optimized method, we have screened a collection of 183 isolates, including 147 received at the Spanish Mycology Reference Laboratory in 2022 and 2023. We have used the ECOFFinder program [39] to calculate the threshold values for non-wild-type behavior, which has permitted the classification of the isolates. In addition, we observed two different subpopulations in the non-wild-type strains, which we tentatively classified as persister and tolerant. To validate this classification, we developed a microscopic assay to analyze the killing dynamics of representative strains. The microscopic method revealed three distinct kinetic profiles that aligned with the previous classification, thereby supporting the validity of the the single-CFU assay. However, we noticed that the killing curves of some persister isolates overlapped with those of susceptible strains, and subsequently we found that persistence did not negatively impact treatment efficacy in our *Galleria m*odel. All together, these results question the biological and clinical relevance of the persistence phenotype detected in our analyses. Future studies should address this issue by (1) elucidating the molecular mechanisms underlying tolerance and persistence, and (2) assessing their relevance in more complex infection models and clinical samples.

Besides, it is surprising that we have detected more non-wild-type strains (tolerant/persister) to IVZ than to VCZ, since the fungicidal capacity of these two antifungals has been demonstrated to be similar [43, 44]. We hypothesize that this may be an *in vitro a*rtifact, like the usual one dilution higher MICs for IVZ than for VCZ. This will need to be addressed once the underlying mechanisms for tolerance and persistence are defined.

Interestingly, we have found that in most occasions tolerance, but not persistence, significantly reduced the therapeutic effect of voriconazole treatment in the *Galleria mellonella i*nfection model. We consider this a highly significant finding, warranting further research to elucidate the potential implication of azole tolerance in therapeutic failure. Furthermore, even if the *Galleria m*odel is useful for preliminary virulence and antifungal screenings, it cannot replicate the intricacies of human infection, such as drug metabolism and pharmacokinetics, interactions of drugs with immune cells or chronic pulmonary pathology [45, 46]. Therefore, more complex models should be used to confirm the actual importance of tolerance and persistence, as they might be underestimated in the *Galleria m*odel.

Altogether, although more research is needed to definitively confirm its impact on treatment failure, based on our results we propose that tolerance should be monitored in clinical isolates and potentially considered to evaluate alternative clinical therapy.

## MATERIAL AND METHODS

### Strains and Media

Two collections of *Aspergillus fumigatus s*trains were used in this study. The first one contains all susceptible isolates (according to the EUCAST E.DEF 9.4) received at the Spanish Mycology Reference Laboratory in 2022 and 2023 (Table S1). The second one contains isolates from environmental origin previously used in persistence studies in our laboratory [34]. All isolates were routinely grown on potato dextrose agar (PDA, Oxoid) for spore harvesting. Spores were always harvested on the same day of MIC or tolerance/persistence determination. For these assays, RPMI-1640 with L-glutamine w/o bicarbonate (Sigma-Aldrich), supplemented with 2% glucose and 0.165 mol/L 3-(N-morpholino) propanesulfonic acid (MOPS, Sigma- Aldrich) was employed.

### MIC determination

The determination of broth dilution minimum inhibitory concentrations of antifungal agents for all *A. fumigatus s*trains was performed following EUCAST reference method E.DEF 9.4 (available at https://www.eucast.org/ast_of_fungi).

### Killing Curves determined by CFUs

An inoculum of 1.25 × 10^5^ conidia/mL of each strain was added to RPMI-1640 containing 8 µg/mL VCZ, 16 µg/mL AMB or 8 µg/mL CSP. 200 µL of the inoculated media were aliquoted in 96-well plates and incubated at 37°C. The full well content was plated on PDA at 0h, 24h or 48h and CFUs were counted after 24 h of incubation at 37°C.

### Establishment of threshold values for tolerance and persistence

To determine the threshold for non-wild-type survival behavior, we employed the ECOFFinder spreadsheet calculator (available at https://clsi.org/resources/ecoffinder/) [39]. To mimic the distribution of strains within a log2 scale (as used in MIC determination), we categorized the strains into groups based on increments of 50 in the number of CFUs obtained after the incubation.

### Single CFU measurement method to detect azole tolerance and persistence

For each strain, a suspension of 2.5×10^6^ conidia/mL was prepared in water and 100 µL inoculated in 100 µL RPMI-1640 2% glucose (see above) containing 8 µg/mL VCZ or IVZ in 96-well plates (2.5×10^5^ conidia/well). Plates were incubated at 35°C for 72 h. After incubation, the plates were centrifuged at 3500 *×g f*or 5 minutes, the drug containing media removed and the wells resuspended in 200 µL of NaCl 0.5M/Tween-20 0.002%. Subsequently, the full content of the wells (200 µL) was inoculated on PDA plates and evenly spread on the surface using a Digralksy loop. PDA plates were incubated at 35°C for ∼30-40 h until colonies were visible. The number of CFUs was counted and related to the initial inoculum to calculate the percentage of killing.

### Killing Curves determined by microscopy

A total of 2.25 x 10^3^ conidia per strain were inoculated in 100 µL of RPMI containing 8 µg/mL VCZ or IVZ and 10 mg/L of SYTOX Red (ThermoFisher Scientific) in a 384 well plate (PhenoPlate TM 384) and incubated at 37°C for 5 h before starting imaging (as virtually no killing is observed in the first 6 h of incubation).

### Image Acquisition

Time-lapse imaging was performed using the Operetta Cell Explorer high-content imaging platform (Revvity). Cells were imaged with a 20× air objective (NA 0.4). For each well, digital phase contrast (DPC) and Sytox fluorescence channels were acquired at 9 positions per well every 15 minutes throughout the experiment. Image capture and instrument control were conducted using Harmony software, version 5.2 (Revvity).

### Image Analysis

Image analysis workflows were developed in Harmony 5.2. Individual cells were identified and segmented in the DPC channel, and cell trajectories were tracked over time. Sytox fluorescence intensity was extracted per cell at each timepoint to determine the onset of membrane-impermeability associated with cell death.

### Computational Classification and Quantification

A custom RStudio script (R v3.6.0 and RStudio v2025.09.2+418; Posit Software, PBC, Boston, MA; *RStudio: Integrated Development Environment for R;* https://www.posit.co/) was developed to classify Sytox-positive (dead) cells over time based on fluorescence thresholds derived from control conditions. For each well, the script calculated (1) the accumulated number of dead cells (cumulative sum across time), and (2) the percentage of dead cells relative to the total number of tracked cells.

### Galleria mellonella infection

Sixth-stage instar larval *G. mellonella m*oths were purchased from Artroposfera (Torrijos, Toledo, Spain). Randomly selected groups of 25 larvae (250 to 4001mg) were injected in the last left proleg with 10 μL of a conidial suspension containing 10^8^ conidia/mL (=10^6^ conidia/larva) of the correspondent *A. fumigatus i*solate using Braun

Omnican 50-U 100 0.5-mL insulin syringes with integrated needles. For relevant groups 10 μL of voriconazole 800 μg/mL was injected in the last right proleg (4 μg/larvae). An untouched control was used to monitor the health of each larvae batch. A PBS control group was subjected to the same treatment, but without fungal infection, to exclude other causes of death beyond *A. fumigatus i*nfection. Survival was followed daily for a period of 7 days. The experiment was repeated two to four independent times. Statistical significance was calculated using the Log-rank (Mantel–Cox) test using GraphPad Prism 10 (GraphPad Software). A *p-*value <0.05 was considered statistically significant.

## ACKNOWLEDGEMENTS

We would like to thank all members of the Spanish Mycology Reference Laboratory for help and support. We acknowledge the use of the microscopes of Advanced Optical Microscopy Unit at ISCIII.

This research has been funded by the AESI (FIS) project PI22CIII/00053.

## CONFLICT OF INTEREST STATEMENT

Ana Alastruey-Izquierdo received funding from Scynexis in the last 36 months to test the preclinical activity of a new antifungal, honoraria from Gilead, Pfizer and Mundipharma for educational talks and participated in an advisory board for Basilea.

The other authors declare no conflict of interest.

## SUPPLEMENTARY MATERIAL

**Table S1.**

List of strains used in this study.

**Table S2.**

Value of the inverse of the slope (1/slope) of the initial 10 hours of incubation for the selected strains.

**Figure S1.**

Itraconazole forms crystals at high concentration (8 µg/mL) and conidia can be seen germinating and growing in their proximity. Two strains are shown **A)** ATCC and **B)** Af293.

