## Supplementary figures and images for "Establishing thresholds for azole tolerance and persistence in *Aspergillus fumigatus t*o study their impact on voriconazole treatment *in vivo*"

### Figure S1

Figure S1

A

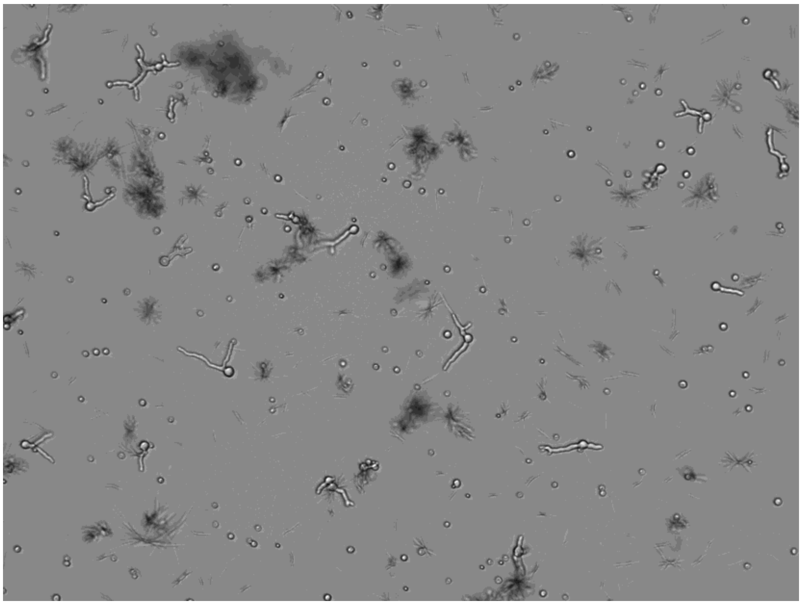

B

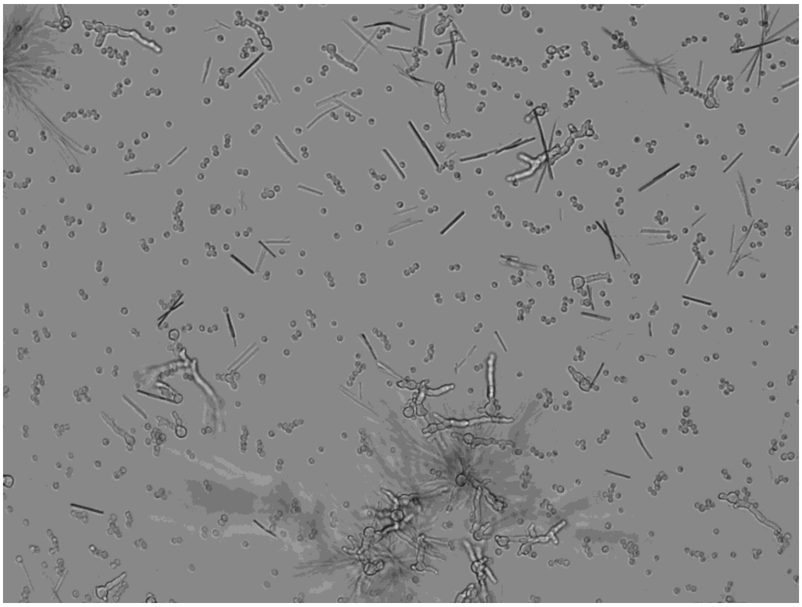
