## Supplementary material for "Establishing thresholds for azole tolerance and persistence in *Aspergillus fumigatus t*o study their impact on voriconazole treatment *in vivo*": Table S1

| **Name** | **Genotype** | **Origin** | **Origin Country** | **Year** |
| --- | --- | --- | --- | --- |
| ATCC 46645 | Wildtype | Clinical | Germany |  |
| Af293 | Wildtype | Clinical | United Kingdom |  |
| MB16 | Wildtype | Enviromental | United Kingdom |  |
| MB28 | Wildtype | Enviromental | United Kingdom |  |
| MB32 | Wildtype | Enviromental | United Kingdom |  |
| MB40 | Wildtype | Enviromental | United Kingdom |  |
| MB49 | Wildtype | Enviromental | United Kingdom |  |
| MB51 | Wildtype | Enviromental | United Kingdom |  |
| MB58 | Wildtype | Enviromental | United Kingdom |  |
| MB59 | Wildtype | Enviromental | United Kingdom |  |
| MB70 | Wildtype | Enviromental | United Kingdom |  |
| MB84 | Wildtype | Enviromental | United Kingdom |  |
| MB105 | Wildtype | Enviromental | United Kingdom |  |
| MB116 | Wildtype | Enviromental | United Kingdom |  |
| MB117 | Wildtype | Enviromental | United Kingdom |  |
| MB125 | Wildtype | Enviromental | United Kingdom |  |
| MB139 | Wildtype | Enviromental | United Kingdom |  |
| MB146 | Wildtype | Enviromental | United Kingdom |  |
| MB149 | Wildtype | Enviromental | United Kingdom |  |
| MB153 | Wildtype | Enviromental | United Kingdom |  |
| PD4 | Wildtype | Clinical | United Kingdom |  |
| PD6 | Wildtype | Clinical | USA |  |
| PD7 | Wildtype | Enviromental | Sweden |  |
| PD8 | Wildtype | Clinical | USA |  |
| PD9 | Wildtype | Enviromental | USA |  |
| PD50 | Wildtype | Enviromental | Ireland |  |
| PD60 | Wildtype | Enviromental | Ireland |  |
| PD104 | Wildtype | Enviromental | Thailand |  |
| PD107 | Wildtype | Enviromental | USA |  |
| PD154 | Wildtype | Enviromental | United Kingdom |  |
| PD249 | Wildtype | Clinical | United Kingdom |  |
| PD251 | Wildtype | Clinical | United Kingdom |  |
| PD254 | Wildtype | Clinical | United Kingdom |  |
| PD259 | Wildtype | Clinical | United Kingdom |  |
| PD264 | Wildtype | Clinical | USA |  |
| PD266 | Wildtype | Clinical | USA |  |
| PD291 | Wildtype | Enviromental | Thailand |  |
| PD292 | Wildtype | Enviromental | Turkey |  |
| CM-10565 | Wildtype | Clinical | Spain | 2022 |
| CM-10571 | Wildtype | Clinical | Spain | 2022 |
| CM-10578 | Wildtype | Clinical | Spain | 2022 |
| CM-10580 | Wildtype | Clinical | Spain | 2022 |
| CM-10585 | Wildtype | Clinical | Spain | 2022 |
| CM-10591 | Wildtype | Clinical | Spain | 2022 |
| CM-10593 | Wildtype | Clinical | Spain | 2022 |
| CM-10594 | Wildtype | Clinical | Spain | 2022 |
| CM-10595 | Wildtype | Clinical | Spain | 2022 |
| CM-10597 | Wildtype | Clinical | Spain | 2022 |
| CM-10598 | Wildtype | Clinical | Spain | 2022 |
| CM-10599 | Wildtype | Clinical | Spain | 2022 |
| CM-10604 | Wildtype | Clinical | Spain | 2022 |
| CM-10607 | Wildtype | Clinical | Spain | 2022 |
| CM-10620 | Wildtype | Clinical | Spain | 2022 |
| CM-10627 | Wildtype | Clinical | Spain | 2022 |
| CM-10633 | Wildtype | Clinical | Spain | 2022 |
| CM-10651 | Wildtype | Clinical | Spain | 2022 |
| CM-10668 | Wildtype | Clinical | Spain | 2022 |
| CM-10675 | Wildtype | Clinical | Spain | 2022 |
| CM-10683 | Wildtype | Clinical | Spain | 2022 |
| CM-10677 | Wildtype | Clinical | Spain | 2022 |
| CM-10686 | Wildtype | Clinical | Spain | 2022 |
| CM-10687 | Wildtype | Clinical | Spain | 2022 |
| CM-10697 | Wildtype | Clinical | Spain | 2022 |
| CM-10699 | Wildtype | Clinical | Spain | 2022 |
| CM-10708 | Wildtype | Clinical | Spain | 2022 |
| CM-10714 | Wildtype | Clinical | Spain | 2022 |
| CM-10716 | Wildtype | Clinical | Spain | 2022 |
| CM-10718 | Wildtype | Clinical | Spain | 2022 |
| CM-10722 | Wildtype | Clinical | Spain | 2022 |
| CM-10723 | Wildtype | Clinical | Spain | 2022 |
| CM-10724 | Wildtype | Clinical | Spain | 2022 |
| CM-10725 | Wildtype | Clinical | Spain | 2022 |
| CM-10756 | Wildtype | Clinical | Spain | 2022 |
| CM-10726 | Wildtype | Clinical | Spain | 2022 |
| CM-10732 | Wildtype | Clinical | Spain | 2022 |
| CM-10733 | Wildtype | Clinical | Spain | 2022 |
| CM-10734 | Wildtype | Clinical | Spain | 2022 |
| CM-10735 | Wildtype | Clinical | Spain | 2022 |
| CM-10743 | Wildtype | Clinical | Spain | 2022 |
| CM-10744 | Wildtype | Clinical | Spain | 2022 |
| CM-10767 | Wildtype | Clinical | Spain | 2022 |
| CM-10765 | Wildtype | Clinical | Spain | 2022 |
| CM-10766 | Wildtype | Clinical | Spain | 2022 |
| CM-10772 | Wildtype | Clinical | Spain | 2022 |
| CM-10773 | Wildtype | Clinical | Spain | 2022 |
| CM-10774 | Wildtype | Clinical | Spain | 2022 |
| CM-10779 | Wildtype | Clinical | Spain | 2022 |
| CM-10781 | Wildtype | Clinical | Spain | 2022 |
| CM-10782 | Wildtype | Clinical | Spain | 2022 |
| CM-10784 | Wildtype | Clinical | Spain | 2022 |
| CM-10794 | Wildtype | Clinical | Spain | 2022 |
| CM-10795 | Wildtype | Clinical | Spain | 2022 |
| CM-10796 | Wildtype | Clinical | Spain | 2022 |
| CM-10822 | Wildtype | Clinical | Spain | 2022 |
| CM-10823 | Wildtype | Clinical | Spain | 2022 |
| CM-10814 | Wildtype | Clinical | Spain | 2022 |
| CM-10861 | Wildtype | Clinical | Spain | 2022 |
| CM-10833 | Wildtype | Clinical | Spain | 2022 |
| CM-10866 | Wildtype | Clinical | Spain | 2022 |
| CM-10869 | Wildtype | Clinical | Spain | 2022 |
| CM-10872 | Wildtype | Clinical | Spain | 2022 |
| CM-10878 | Wildtype | Clinical | Spain | 2022 |
| CM-10881 | Wildtype | Clinical | Spain | 2022 |
| CM-10884 | Wildtype | Clinical | Spain | 2022 |
| CM-10891 | Wildtype | Clinical | Spain | 2022 |
| CM-10892 | Wildtype | Clinical | Spain | 2022 |
| CM-10894 | Wildtype | Clinical | Spain | 2022 |
| CM-10907 | Wildtype | Clinical | Spain | 2022 |
| CM-10909 | Wildtype | Clinical | Spain | 2022 |
| CM-10911 | Wildtype | Clinical | Spain | 2022 |
| CM-10898 | Wildtype | Clinical | Spain | 2022 |
| CM-10919 | Wildtype | Clinical | Spain | 2022 |
| CM-10929 | Wildtype | Clinical | Spain | 2022 |
| CM-10930 | Wildtype | Clinical | Spain | 2022 |
| CM-10934 | Wildtype | Clinical | Spain | 2022 |
| CM-10939 | Wildtype | Clinical | Spain | 2022 |
| CM-10940 | Wildtype | Clinical | Spain | 2022 |
| CM-10966 | Wildtype | Clinical | Spain | 2022 |
| CM-10954 | Wildtype | Clinical | Spain | 2022 |
| CM-11010 | Wildtype | Clinical | Spain | 2022 |
| CM-11024 | Wildtype | Clinical | Spain | 2022 |
| CM-10972 | Wildtype | Clinical | Spain | 2022 |
| CM-11027 | Wildtype | Clinical | Spain | 2022 |
| CM-10979 | Wildtype | Clinical | Spain | 2022 |
| CM-11049 | Wildtype | Clinical | Spain | 2022 |
| CM-10991 | Wildtype | Clinical | Spain | 2022 |
| CM-11018 | Wildtype | Clinical | Spain | 2022 |
| CM-11038 | Wildtype | Clinical | Spain | 2022 |
| CM-11063 | Wildtype | Clinical | Spain | 2022 |
| CM-11066 | Wildtype | Clinical | Spain | 2022 |
| CM-11048 | Wildtype | Clinical | Spain | 2023 |
| CM-11074 | Wildtype | Clinical | Spain | 2023 |
| CM-11075 | Wildtype | Clinical | Spain | 2023 |
| CM-11077 | Wildtype | Clinical | Spain | 2023 |
| CM-11078 | Wildtype | Clinical | Spain | 2023 |
| CM-11090 | Wildtype | Clinical | Spain | 2023 |
| CM-11098 | Wildtype | Clinical | Spain | 2023 |
| CM-11105 | Wildtype | Clinical | Spain | 2023 |
| CM-11114 | Wildtype | Clinical | Spain | 2023 |
| CM-11121 | Wildtype | Clinical | Spain | 2023 |
| CM-11128 | Wildtype | Clinical | Spain | 2023 |
| CM-11132 | Wildtype | Clinical | Spain | 2023 |
| CM-11134 | Wildtype | Clinical | Spain | 2023 |
| CM-11137 | Wildtype | Clinical | Spain | 2023 |
| CM-11138 | Wildtype | Clinical | Spain | 2023 |
| CM-11146 | Wildtype | Clinical | Spain | 2023 |
| CM-11170 | Wildtype | Clinical | Spain | 2023 |
| CM-11172 | Wildtype | Clinical | Spain | 2023 |
| CM-11174 | Wildtype | Clinical | Spain | 2023 |
| CM-11185 | Wildtype | Clinical | Spain | 2023 |
| CM-11186 | Wildtype | Clinical | Spain | 2023 |
| CM-11189 | Wildtype | Clinical | Spain | 2023 |
| CM-11207 | Wildtype | Clinical | Spain | 2023 |
| CM-11202 | Wildtype | Clinical | Spain | 2023 |
| CM-11205 | Wildtype | Clinical | Spain | 2023 |
| CM-11216 | Wildtype | Clinical | Spain | 2023 |
| CM-11228 | Wildtype | Clinical | Spain | 2023 |
| CM-11241 | Wildtype | Clinical | Spain | 2023 |
| CM-11245 | Wildtype | Clinical | Spain | 2023 |
| CM-11252 | Wildtype | Clinical | Spain | 2023 |
| CM-11268 | Wildtype | Clinical | Spain | 2023 |
| CM-11285 | Wildtype | Clinical | Spain | 2023 |
| CM-11288 | Wildtype | Clinical | Spain | 2023 |
| CM-11292 | Wildtype | Clinical | Spain | 2023 |
| CM-11294 | Wildtype | Clinical | Spain | 2023 |
| CM-11311 | Wildtype | Clinical | Spain | 2023 |
| CM-11337 | Wildtype | Clinical | Spain | 2023 |
| CM-11352 | Wildtype | Clinical | Spain | 2023 |
| CM-11363 | Wildtype | Clinical | Spain | 2023 |
| CM-11393 | Wildtype | Clinical | Spain | 2023 |
| CM-11409 | Wildtype | Clinical | Spain | 2023 |
| CM-11418 | Wildtype | Clinical | Spain | 2023 |
| CM-11427 | Wildtype | Clinical | Spain | 2023 |
| CM-11429 | Wildtype | Clinical | Spain | 2023 |
| CM-11444 | Wildtype | Clinical | Spain | 2023 |
| CM-11471 | Wildtype | Clinical | Spain | 2023 |
| CM-11489 | Wildtype | Clinical | Spain | 2023 |
| CM-11498 | Wildtype | Clinical | Spain | 2023 |
| CM-11525 | Wildtype | Clinical | Spain | 2023 |
| CM-11528 | Wildtype | Clinical | Spain | 2023 |
| CM-11533 | Wildtype | Clinical | Spain | 2023 |
| CM-11542 | Wildtype | Clinical | Spain | 2023 |
| CM-11544 | Wildtype | Clinical | Spain | 2023 |
| CM-11547 | Wildtype | Clinical | Spain | 2023 |
| CM-11548 | Wildtype | Clinical | Spain | 2023 |
