## Supplementary material for "Establishing thresholds for azole tolerance and persistence in *Aspergillus fumigatus t*o study their impact on voriconazole treatment *in vivo*": Table S2

| Strain | Survival Rate (1/slope) first 8 h | |
| --- | --- | --- |
|  | VZC | IVZ |
| ATCC | 1.219 | 1.148 |
| CM-10565 | 1.589 | 1.578 |
| CM-10773 | 1.746 | 1.883 |
| CM-11134 | 2.083 | 2.221 |
| CM-10597 | 4.882 | 6.256 |
| CM-10779 | 4.208 | 4.337 |
| CM-10929 | 5.249 | 4.344 |
| CM-10881 | 3.264 | 3.202 |
| CM-11429 | 2.945 | 3.409 |
| Af293 | 9.915 | 10.57 |
| CM-10571 | 28.35 | 21.31 |
| CM-10677 | 31.95 | 31.45 |
| CM-11185 | 10.11 | 10.79 |
| CM-11189 | 50.17 | 42.45 |
